# Inositol lipid synthesis is widespread in host-associated Bacteroidetes

**DOI:** 10.1101/2021.04.26.441525

**Authors:** S. L. Heaver, H. H. Le, P. Tang, A. Baslé, J. Marles-Wright, E. L. Johnson, D. J. Campopiano, R. E. Ley

## Abstract

Ubiquitous in eukaryotes, inositol lipids have finely tuned roles in cellular signaling and membrane homeostasis. In Bacteria, however, inositol lipid production is rare. Recently, the prominent human gut bacterium *Bacteroides thetaiotaomicron* (BT) was reported to produce inositol lipids, including inositol sphingolipids, but the pathways remain ambiguous and their prevalence unclear. Here, we investigated the gene cluster responsible for inositol lipid synthesis in BT using a novel strain with inducible control of sphingolipid synthesis. We characterized the biosynthetic pathway from *myo-*inositol-phosphate (MIP) synthesis to phosphoinositol-dihydroceramide, including structural and kinetic studies of the enzyme MIP synthase (MIPS). We determined the crystal structure of recombinant BT MIPS with bound NAD cofactor at 2.0 Å resolution, and identified the first reported phosphatase for the conversion of bacterially-derived phosphatidylinositol phosphate (PIP) to phosphatidylinositol (PI). Transcriptomic analysis indicated inositol production is nonessential but its loss alters BT capsule expression. Bioinformatic and lipidomic comparisons of Bacteroidetes species revealed a novel second putative pathway for bacterial PI synthesis without a PIP intermediate. Our results indicate that inositol sphingolipid production, via one of the two pathways, is widespread in host-associated Bacteroidetes, and may be implicated in host interactions both indirectly via the capsule and directly through inositol lipid provisioning.

## Introduction

Inositol, a carbocyclic sugar abundant in eukaryotes, forms the structural basis for diverse phosphorylated secondary messenger inositol phosphates and inositol lipids. At their simplest, inositol lipids have inositol as their polar headgroup, as is the case with phosphatidylinositol (PI; on a glycerophospholipid backbone) or inositol phosphorylceramide (on a sphingolipid backbone; Fig. 1A). The inositol headgroup can be subject to further modifications, including the phosphorylation of PI at multiple sites to form bioactive phosphoinositides, or the addition of a mannose on inositol sphingolipids to form the mannosylinositol phosphorylceramides (MIPCs) abundant in yeast ^1,2^. Inositol derivatives control key processes of eukaryotic cell physiology. For instance, although phosphoinositides constitute a small fraction of overall phospholipids, they are ubiquitous and essential in roles such as marking organelle identity, regulating cytoskeleton-membrane interactions, and controlling cell division and autophagy ^3,4^.

**Figure 1.**
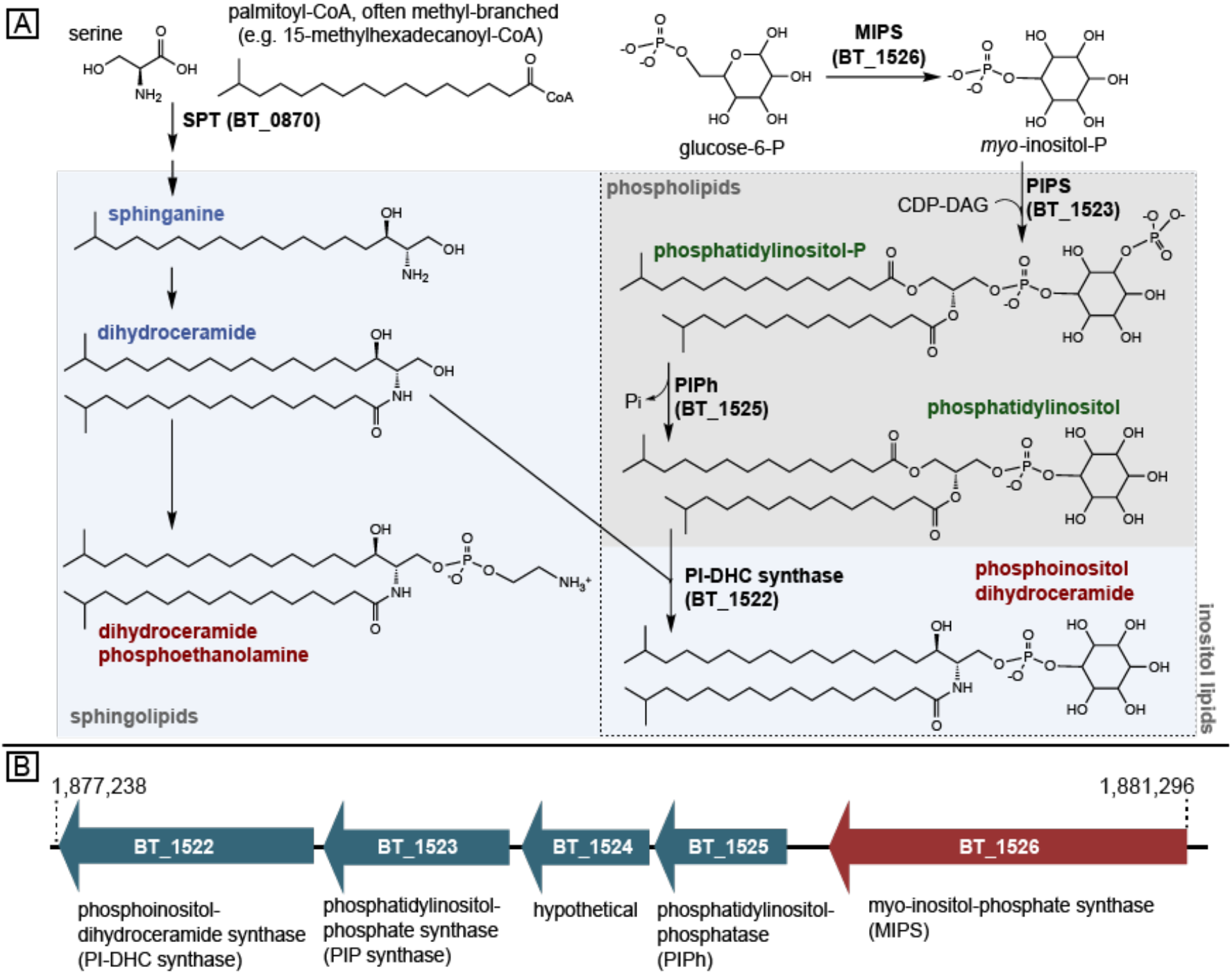
Enzymatic pathway for inositol lipid synthesis in BT. (A) The *de novo* sphingolipid synthesis metabolic pathway in relation to inositol lipid synthesis, with BT enzymes investigated in this study (bolded black) and representative lipid structures. Sphingolipid structures are on a light blue background; phospholipid structures are on a gray background. (B) The genomic region of BT inositol and inositol lipid synthesis. Gene color (blue or red) indicates membership in an operon predicted by BioCyc. Annotations are of the enzyme functions elucidated in this study (due to the lack of a lipid phenotype in its knockout strain, BT_1524 was not investigated further and remains hypothetical).

Despite the widespread distribution of inositol lipids in eukaryotes, relatively little is known about the structure and distribution of inositol lipids in Bacteria. Bacterial inositol lipids are comparatively rare across bacterial species, with production previously thought to be limited largely to phosphatidylinositol (PI) synthesis in members of the phylum Actinobacteria (*e*.*g*., *Mycobacteria, Corynebacteria*, and *Streptomyces*) ^5–7^ and Spirochaetes (*Treponema*) ^8^. In *Mycobacterium tuberculosis*, an obligate intracellular human pathogen, the inositol headgroups of PI are the outer membrane molecules to which surface oligosaccharide virulence factors are linked ^9^.

In eukaryotes, both sphingolipids (SLs, lipids with a sphingosine, or long-chain base backbone) and inositol lipids are involved in the regulation of cell fate and differentiation, inflammation, protein trafficking, and gene regulation in central metabolic pathways, with imbalances linked with the pathologies of a growing inventory of diseases ^4,10–12^. These two lipid types intersect in the inositol sphingolipids, such as the glycosylinositol phosphorylceramides abundant in yeast and plants. Inositol SLs are known to be produced by the periodontal pathogen *Tannerella forsythia* ^13^, a member of the Bacteroidetes. Furthermore, Brown *et al*. recently reported inositol SLs in the common human gut commensal *Bacteroides thetaiotaomicron* ^*14*^. Ceramide phosphoryl-*myo*-inositol has been reported in *Sphingobacterium spiritivorum* ^15^, a free-living member of the Bacteroidetes, and more recently in *Myxococcus xanthus* ^16^, of the Proteobacteria phylum. In contrast to the well-studied inositol lipids, including inositol SLs, in plants and fungi, bacterial inositol sphingolipid synthesis has been largely overlooked.

The discovery of inositol lipids in a several more species suggests that these lipids may be more widespread in bacteria than previously thought and may use novel pathways. Across kingdoms, *de novo* inositol synthesis begins with the formation of inositol phosphate from glucose 6-phosphate (G6P) by a *myo*-inositol phosphate synthase (MIPS, EC 5.5.1.4) ^11^. From here, a bacterial pathway for inositol glycerophospholipid synthesis is mostly known (*e*.*g*., in *Mycobacteria*), and differs from the eukaryotic pathway by the direct use of inositol-phosphate, not its dephosphorylated inositol form, as a substrate in the formation of PI. This leads first to the synthesis of phosphatidylinositol-phosphate (PIP) from CDP-diacylglycerol (CDP-DAG) and MIP, which is subsequently dephosphorylated to PI ^17^. Though the PIP synthase has been well characterized in bacteria ^18^, the phosphatase responsible for the conversion of bacterial PIP to PI has not yet been identified ^18^. In addition, though the gene cluster for bacterial inositol SL synthesis has been predicted in *B. thetaiotaomicron* (hereafter BT) ^14^, the functions of these enzymes remain to be confirmed.

Here, we combine genomic and biochemical approaches to functionally characterize the predicted inositol lipid metabolism gene cluster in BT from the initial synthesis of *myo-*inositol-phosphate (MIP) to its addition as a headgroup to glycerophospholipids and SLs. Together with the description of a novel putative alternative gene cluster, common in the *Prevotella*, this work broadens the understanding of how gut bacteria synthesize complex lipids, and reveals an extensive capacity for inositol lipid synthesis among gut-associated Bacteroidetes.

## Results and Discussion

We first identified genes responsible for inositol lipid metabolism in BT (Fig. 1A). We identified BT_1522 by NCBI Blast-P as having high homology to the yeast enzyme inositol phosphorylceramide synthase (IPC synthase, also known as AUR1) that catalyzes the attachment of the phophorylinositol group onto ceramide (query cover 50%, e-value 1e-15, percent identity 26%). This led us to hypothesize that this enzyme is responsible for phosphoinositol SL synthesis in BT, though BT SLs have predominantly dihydroceramide (not ceramide) backbones, leading instead to the synthesis of phosphoinositol dihydroceramide (PI-DHC). *BT_1522* and its gene cluster (Fig. 1B) were previously predicted to be involved in inositol lipid metabolism ^14^. Adjacent predicted genes in the cluster include BT_1523 (annotated as a CDP-diacylglycerol-inositol 3-phosphatidyltransferase), BT_1524 (hypothetical protein), BT_1525 (currently annotated as phosphatidylglycerophosphatase A, PgpA), and BT_1526 (*myo-*inositol phosphate synthase, “MIPS”).

We next constructed a BT strain with tunable SL synthesis. As inositol lipids in BT include both glycerophospholipids (PI) and SLs (PI-DHC), we created a strain of BT with inducible control of the first enzyme in the *de novo* SL synthesis pathway, serine palmitoyltransferase (SPT; BT_0870; Fig. 1) ^19^. This inducible-SPT (“iSPT”) strain enables precise control over BT synthesis to produce both PI and PI-DHC, or solely PI. As expected, in the absence of SPT, we detected no SLs by thin-layer chromatography (TLC) analysis (Fig. 2A). However, in iSPT, SL synthesis gradually increased with increasing levels of the anhydrotetracycline (aTC) inducer, to an approximate wild-type (WT) SL abundance with 100 ng/mL aTC induction (Fig. 2A). At full induction, SLs composed roughly half of the extracted lipids measured by TLC densitometry (47±7%, n=6). These SLs include PI-DHC and phosphoethanolamine dihydroceramide (PE-DHC), among others (Fig. 2C).

**Figure 2.**
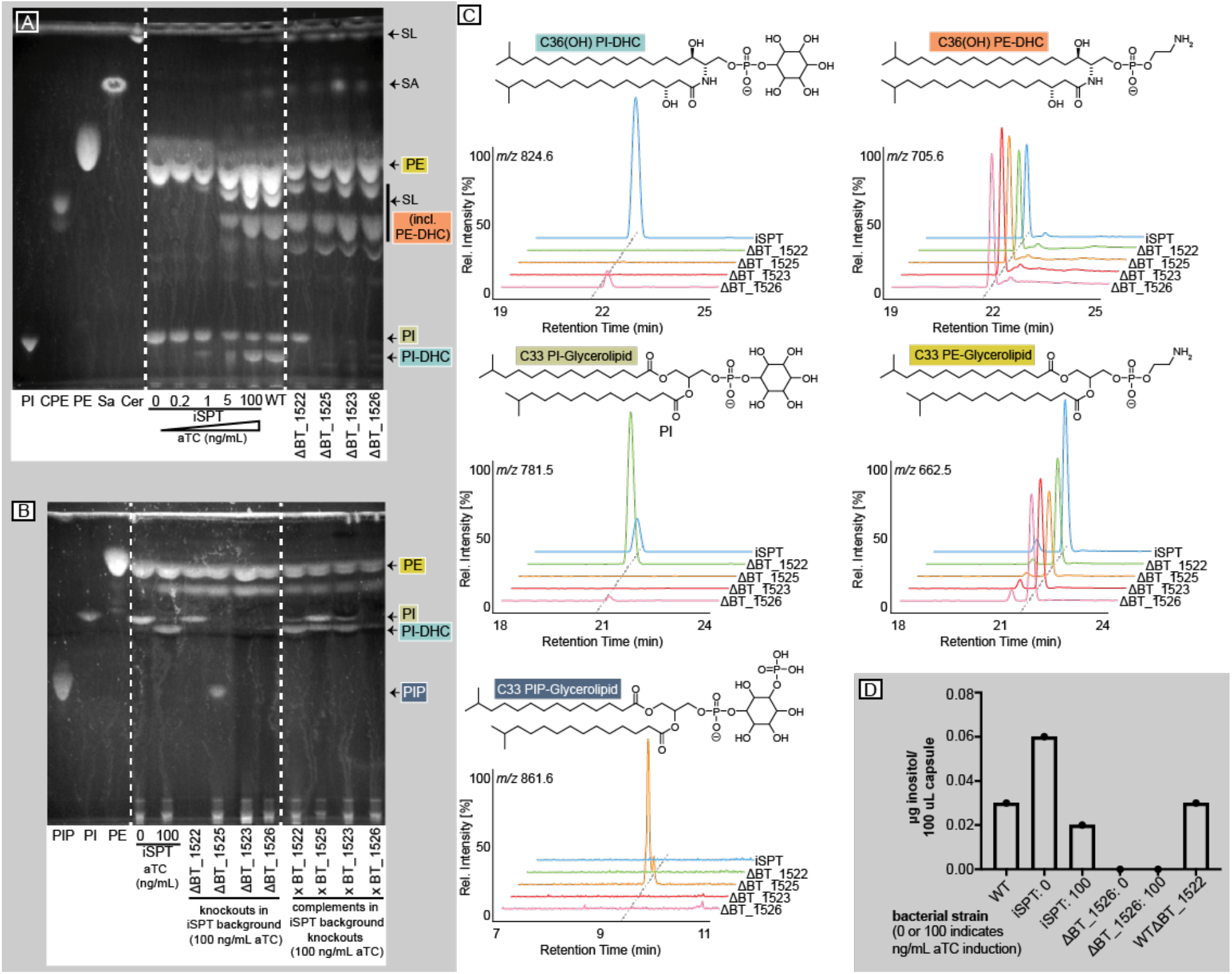
BT produces inositol phospholipids and sphingolipids. (A) Thin layer chromatography (TLC) of left to right: first section indicated by dashed lines includes five standards: PI = 16:0 phosphatidylinositol; PIP = 18:1 PI(3)P; CPE = ceramide phosphoethanolamine; PE = egg yolk phosphatidylethanolamine, Sa = d18:0 sphinganine, Cer = d18:1/18:0 ceramide; second section includes six standard (non-acidic) lipid extracts from the iSPT BT strain (used as a background for knockout generation) at 0, 0.2, 1, 5 and 100 ng/mL aTC induction of SPT, and WT BT VPI-5482; third section includes standard lipid extraction from ΔBT_1522, ΔBT_1523, ΔBT_1525, and ΔBT_1526 knockout strains in the iSPT background at 100 ng/mL aTC induction of SPT. (B) TLC of: standards (PIP, PI, PE) as in panel A; second section shows PIP lipid extractions of iSPT strains at 0 and 100 ng/mL aTC followed by ΔBT_1522, ΔBT_1523, ΔBT_1525, ΔBT_1526, and each of their respective complementations (all strains in the iSPT background) at 100 ng/mL aTC induction of SPT. (C) Predicted structures and ion chromatograms demonstrating detection of inositol lipids and sphingolipids in iSPT, ΔBT_1522, ΔBT_1523, ΔBT_1525, and ΔBT_1526 (all strains in the iSPT background) at 100 ng/mL aTC induction. (D) Quantification of inositol in the capsule of BT strains with or without sphingolipids (iSPT 100 vs. 0 ng/mL aTC induction), PI-DHC synthase (WTΔBT_1522), and MIPS (ΔBT_1526, at 0 and 100 ng/mL aTC induction).

To uncover the function of each predicted enzyme in the putative inositol lipid metabolism pathway, we knocked out the individual genes (BT_1522 to BT_1526) in the iSPT background by scarless deletion ^20^ (denoted ΔBT_1522 to ΔBT_1526). BT_1522 was also knocked out in the WT background (indicated by WTΔBT_1522). We examined the lipid content of the resulting knockout strains (with SL synthesis fully induced) using TLC and HPLC-MS. Consistent with the predicted role for BT_1522 as a PI-DHC synthase, the ΔBT_1522 strain failed to produce PI-DHC, but production of PI and non-inositol SLs, including PE-DHC, was unaltered (Fig. 2A-C). Similarly, the ΔBT_1526 strain (lacking the predicted MIPS) failed to produce both PI and PI-DHC, in accordance with the loss of the *myo*-inositol-phosphate substrate.

Interestingly, both the ΔBT_1523 and ΔBT_1525 strains also failed to produce both PI and PI-DHC (Fig. 2A). As the synthesis of other glycerophospholipids did not appear to be affected in the ΔBT_1525 strain, this observation was not in agreement with the annotated function of BT_1525 as a PgpA ^21^. We hypothesized that BT may use a two-step process to synthesize PI similar to that found in *Mycobacteria*, which uses a PIP intermediate ^18^. In accord, comparison of the functional protein motifs in BT_1523 and BT_1525 with those in the characterized *Renobacterium salmoninarum* PIP synthase (PIPS) ^22^ revealed the same conserved catalytic residues (DX_2_DGX_2_AR…GX_3_DX_3_D) in BT_1523. This observation supports the notion that BT_1523 functions as a PIPS in the biosynthesis of PIP.

To our knowledge, PIP has not been reported in BT. This is likely due, in part, because PIP extractions under non-acidic conditions may be low yielding. Also, PIP abundance may be inherently low in BT, similar to low phosphoinositide abundances in eukaryotes ^3^. Following a PIP-optimized lipid extraction of BT and the knockout strains, we detected high levels of PIP in ΔBT_1525, detectable by both TLC and HPLC-MS (Fig. 2B-C). PIP accumulation in ΔBT_1525 suggests BT_1525 is most likely a phosphatidylinositolphosphatase (PIPh), responsible for the rapid downstream conversion of PIP to PI. Though BT_1525 has homology to the phosphatidylglycerophosphatase A protein family (Pfam), this sequence similarity could reflect an expansion of the functional role of this protein motif from the dephosphorylation of a phospholipid glycerophosphate headgroup to an inositolphosphate headgroup. Previous work has shown that transcriptomic expression of *BT_1525* is higher than *BT_1523* and *BT_1522* in every growth phase of BT ^23^, likely enabling the rapid conversion of PIP to PI and preventing accumulation of PIP in BT.

Despite the central location of *BT_1524* in the inositol lipid metabolism gene cluster, inositol lipids in ΔBT_1524 phenocopied BT by TLC. The *BT_1524* gene is predicted to encode an integral membrane protein with a Gtr-A motif (Pfam); other Gtr-A family proteins are involved in cell surface polysaccharide or exopolysaccharide synthesis ^24–26^, suggesting BT_1524 may be involved in the membrane transfer of an inositol-linked lipid. Due to a lack of a detectable lipid phenotype in this mutant, we did not investigate it further.

To confirm that the loss of inositol lipids in knockout strains was not due to off-target effects, the native BT sequence of each gene was integrated genomically into knockout strains in the iSPT background (ΔBT_1522, ΔBT_1523, ΔBT_1525, ΔBT_1526), paired with a constitutive promoter optimized for BT ^27^. The complementation was successful for three of the four strains (ΔBT_1522, ΔBT_1523, and ΔBT_1525), fully restoring the capacity for both PI and PI-DHC synthesis (Fig. 2B).

To confirm the predicted function of BT_1526 as a redox-neutral, NAD^+^/NADH-dependent MIPS, we cloned the gene and heterologously overexpressed the protein in *E. coli* (Fig. 3 and Supporting Information). The N-terminally His-tagged BT_1526 expressed well in a highly soluble form (∼50 kDa in size, observed by SDS-PAGE and confirmed by electrospray ionization mass spectrometry, see SI), and was purified to homogeneity by standard immobilized metal affinity chromatography methods. The MIPS activity of BT_1526 was confirmed using a colorimetric endpoint assay which monitors the appearance of the inorganic phosphate released from the MIP product and not the G6P substrate ^28^ (Fig. 3A). Kinetic analysis of BT_1526 MIPS operating on G6P are as follows: K_m_ = 9.97 ± 0.94 mM, V_max_ = 26.17 ± 0.10 µM/min, specific activity = 0.513 µmol/min/mg (Fig. 3B). This activity is in the range of the published specific activities of MIPS from *S. cerevisiae* (0.41 µmol/min/mg) ^29,30^, *Synechocystis* sp. (0.02 µmol/min/mg) ^29^, *A. fulgidus* (11.8 µmol/min/mg) ^31^, and *A. thaliana* (∼0.1 µmol/min/mg) ^32^.

**Figure 3.**
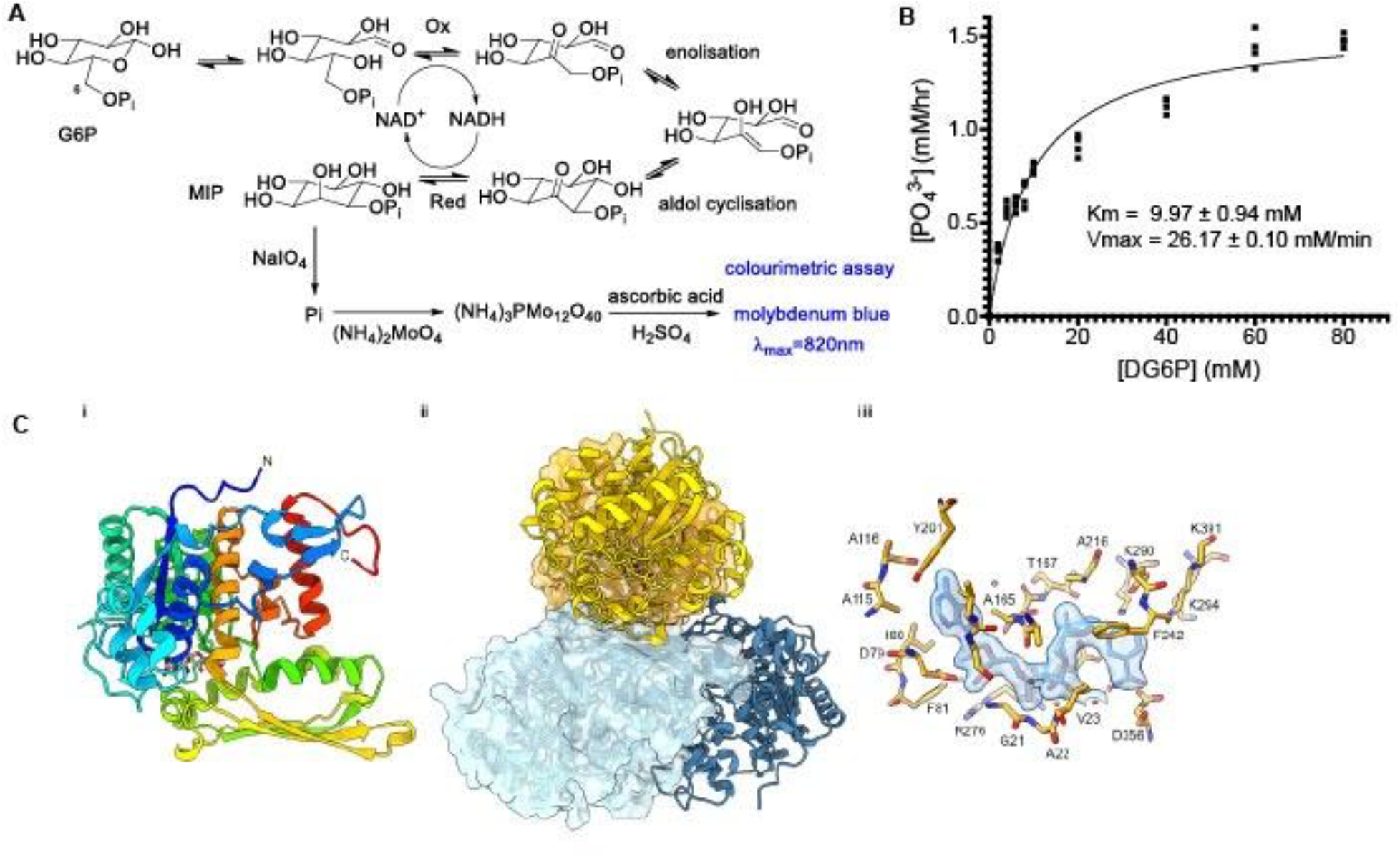
BT_1526 produces *myo*-inositol-phosphate *in vitro*. (A) Proposed mechanism for the MIPS-catalyzed NAD-dependent/redox-neutral conversion of G6P to MIP. (B) Molybdenum blue assay for detection of MIP. Kinetic analysis of recombinant BT_1526 MIPS using G6P as substrate. (C) The crystal structure of BT_1526 MIPS: (i) The monomer subunit, (ii) the tetramer, (iii) the structure of the MIPS:NAD complex.

We further characterized the BT_1526 protein by determining the X -ray crystal structure to 2.0 Å resolution by molecular replacement using a model derived from the *Archaeoglobus fulgidus* MIPS structure (PDBID: 3QVT) (refinement statistics in Supplementary Table 1). The overall 3D fold of BT_1526 is consistent with other members of this family with a Rossman fold-like nucleotide binding domain (residues 4-244, 355-429) with an intercalated catalytic/dimerization domain (residues 245-354) ^33–36^. The protein adopts a tetrameric dimer of dimers quaternary structure of approximately 188 kDa, with the dimerization domain forming an extended beta-sheet between monomers to create a saddle-like interface for the two dimers within the tetramer. Though the protein was purified without the addition of any cofactors or substrates, strong electron density consistent with the NAD^+^ cofactor was observed in the initial maps calculated after molecular replacement and the final structure contains an NAD^+^ molecule associated with each chain modelled at unit occupancy. Given the redox neutrality of the MIPS enzyme (*i*.*e*., it catalyzes substrate oxidation, then reduction) there is a clear benefit to the retention of the NAD^+^ in the active site for the lifetime of the protein, in agreement with NAD^+^ retention in MIPS from other species ^37^. The catalytic active site region of the protein is well conserved among members of the MIPS family for which a structure has been determined, with a cluster of lysine and aspartic acid residues responsible for binding and orienting the G6P substrate for isomerization to the MIP product, highlighting the importance of these residues for the correct activity of the enzyme. Outside of the highly conserved ligand binding site, the overall fold and quaternary structure of the representatives of the family in the PDB is highly conserved, although eukaryotic MIPS proteins have an N-terminal extension which appears to further stabilize the quaternary structure of the protein (Supp. Fig. 4-5). Overall, the enzyme described here as the first MIPS representative in the Bacteroidetes retains the key functional elements of other MIPS enzymes, underscoring its conservation across biological kingdoms.

**Figure 4.**
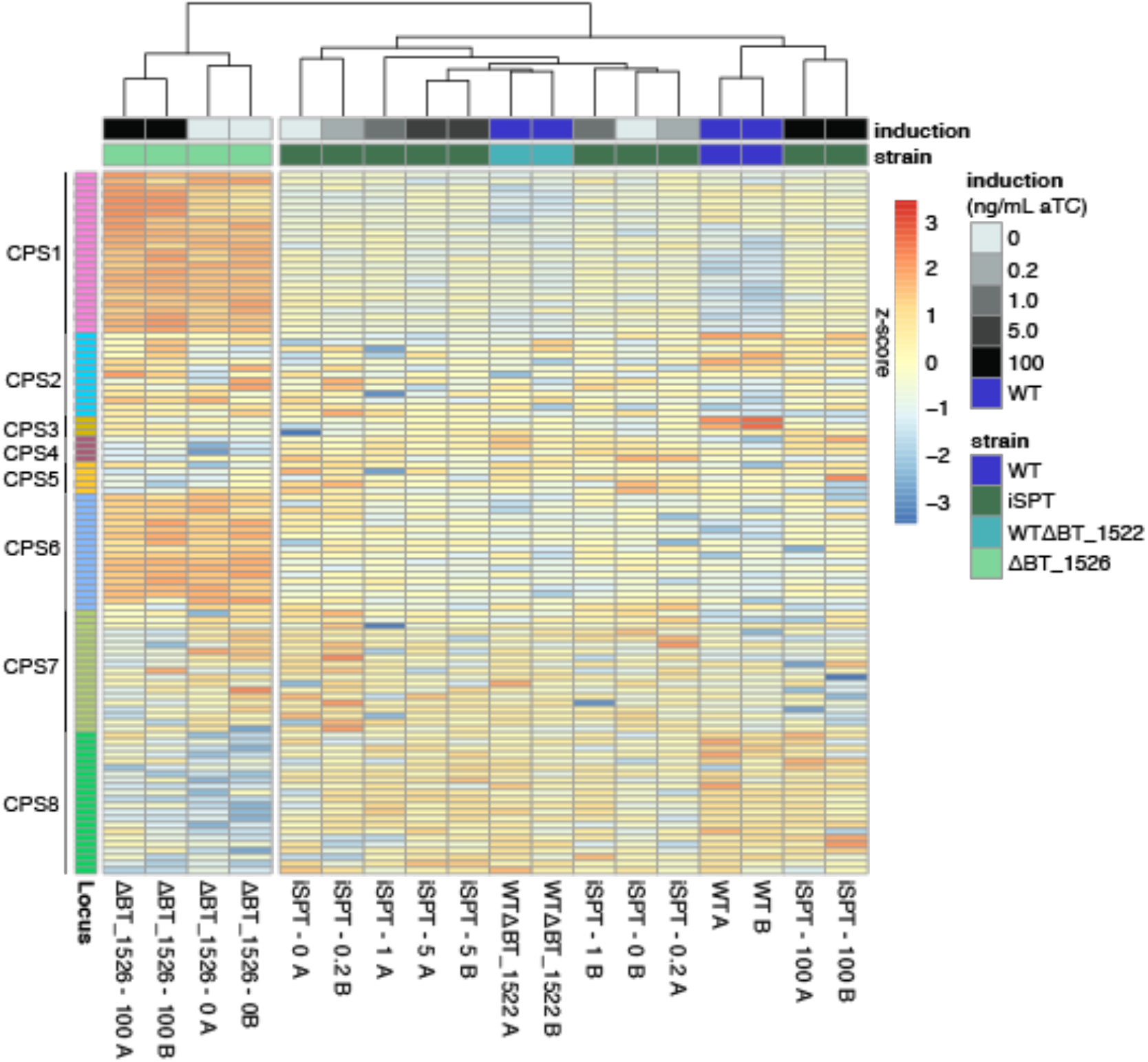
Deletion of MIPS (BT_1526) alters expression of genes for capsular polysaccharide synthesis pathway loci in BT. Gene expression data (normalized log_2_ expression values, scaled by row, with Euclidean column clustering) in the 8 BT capsular polysaccharide synthesis (CPS) loci. For easier visualization, genes were filtered to include those in which maximum log_2_-normalized expression is > 1.5 and exclude those with maximum absolute log_2_-fold-change difference in expression < 1.5 in all pairwise comparisons of conditions. Color in the far left column indicates gene assignment to one of 8 CPS loci. Strains tested include WT BT, iSPT, WTΔBT_1522, and ΔBT_1526 in the iSPT background. SPT induction in the iSPT strains at 0, 0.2, 1.0, 5.0, or 100 ng/mL aTC induction is indicated in shades of grey. Labels below each column indicate strain and aTC induction level (ng/mL aTC) redundantly with the color key; “A” and “B” labels represent biological replicates.

To determine the role of MIPS in BT, we compared gene transcription of the ΔBT_1526 strain to its iSPT strain at full SPT induction. Lacking MIPS, ΔBT_1526 cells lack not only inositol lipids (PIP/PI/PI-DHC), but also any other molecules for which inositol may be used as a substrate, such as cell surface polysaccharides. Only 29 genes were differentially expressed with greater than 1.5 absolute log_2_-fold-change in the ΔBT_1526 strain compared to the background iSPT strain at 100 ng/mL induction of SPT. These genes were almost entirely involved in capsule biosynthesis (Table S3). Expression of CPS loci was fairly uniform across varied levels of SPT induction in the iSPT strain, while the ΔBT_1526 strain had notable upregulation of capsular polysaccharide synthesis loci 1 and 6 (CPS1 and CPS6) (Fig. 4).

To assess whether PI-DHC loss alone was responsible for the capsule effect in BT, we monitored the transcriptional response of WTΔBT_1522 at early stationary phase, grown in minimal medium with glucose as the sole carbon source. Compared to its BT background strain, the WTΔBT_1522 strain had differential expression of 37 genes above a cutoff of 1.5 absolute log_2_-fold-change (Table S2). These included many hypothetical proteins and membrane-associated proteins, including those involved in carbohydrate metabolism, such as SusD starch-binding protein homologs (BT_3025 and BT_2806). Pathway enrichment analysis of these 37 genes showed an enrichment of transcripts involved in sugar degradation (specifically, 5-dehydro-4-deoxy-D-glucuronate degradation, p = 0.006, Fisher exact test, Benjamini-Hochberg correction), acetate and ATP formation from acetyl-CoA (p = 0.098), and carboxylate degradation (p = 0.098). In plants and yeasts, which also produce inositol-linked SLs, these lipids are critical for fundamental aspects of the organism’s physiology, for example, protein anchoring and programmed cell death in plants ^38^. The yeast homolog of BT_1522 is an antifungal target, highlighting its core function in yeast physiology, which is inhibited by the cyclic depsipeptide natural product aureobasidin (hence the name AUR1) ^39^. Though WTΔBT_1522 had few transcriptomic changes relative to WT controls, its affected pathways appear central to carbohydrate degradation and energy synthesis. As such, other inositol derivatives or PI, not PI-DHC, are implicated in the altered capsule expression of the ΔBT_1526 strain.

Inositol has not been previously reported as a component of BT capsule ^40^, perhaps due to its common use as an internal standard in the HPAEC-PAD analysis of capsule components. Using an alternative standard, we detected inositol in the capsular monosaccharide components of WT BT, iSPT, and WTΔBT_1522 strains (Fig. 2D). However, we did not detect inositol in the capsules of SPT-induced and -uninduced ΔBT_1526 strains. To assess whether the transcriptional trend of the ΔBT_1526 to upregulate CPS1 and CPS6 was reflected in the visible cell capsule, we performed scanning electron microscopy on iSPT and ΔBT_1526 strains with and without SL induction (Fig. 5A). Capsule structures were heterogeneous in both strains, but in comparison to the iSPT background strain, more of the cells of the ΔBT_1526 strain exhibited a dense structure extending from the cell surface, with apparent exopolysaccharide connecting adjacent cells. This was particularly noticeable when SL synthesis was not induced.

**Figure 5.**
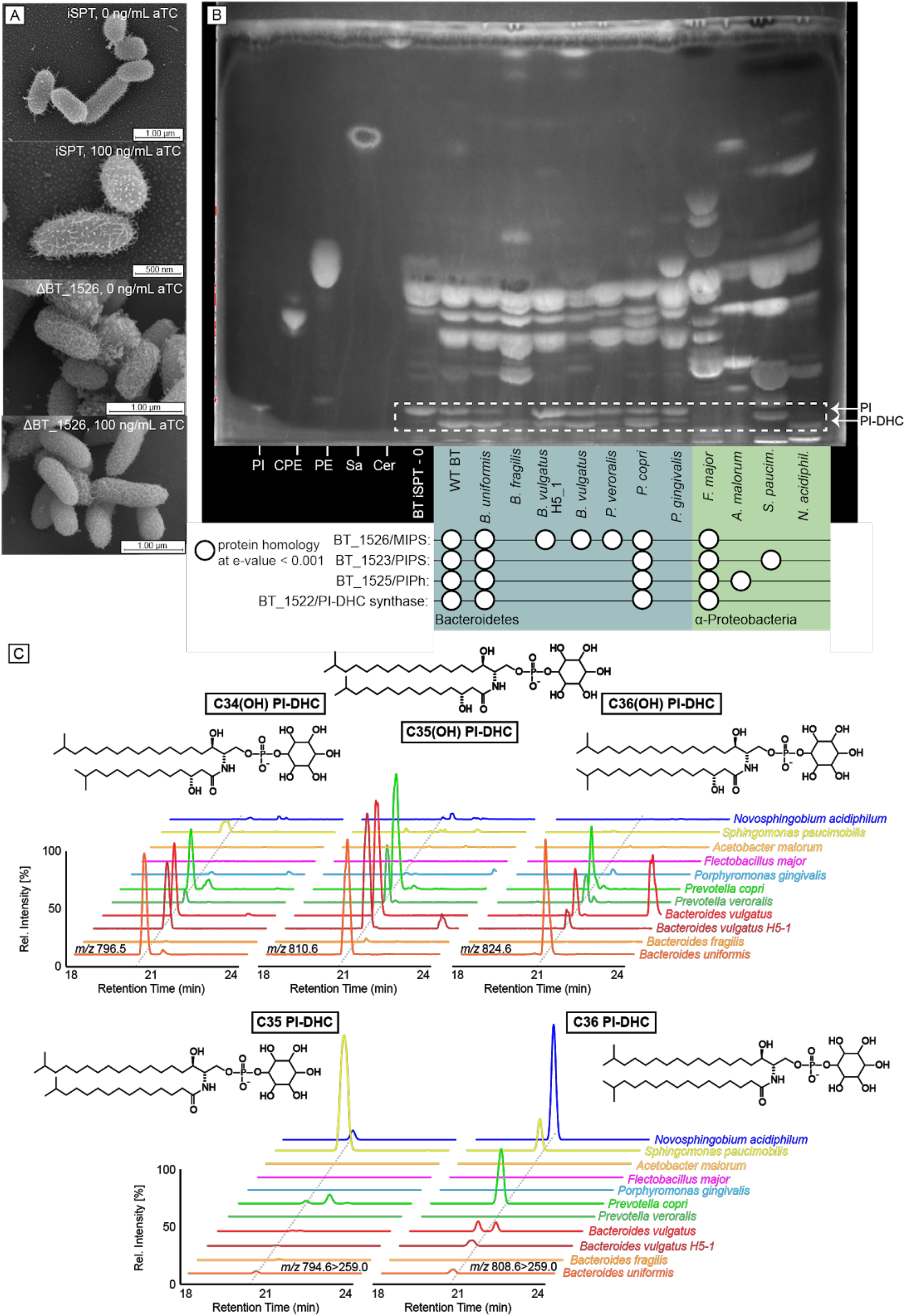
The capacity to produce PI-DHC is widespread among sphingolipid-producing bacteria. (A) Scanning electron microscopy of the iSPT and ΔBT_1526 strains at 0 and 100 ng/mL aTC induction of SPT grown in minimal medium. (B) TLC of lipid standards and lipid extractions from a diverse array of sphingolipid-producing bacteria. Lanes 1-5, left to right: PI = 16:0 phosphatidylinositol; CPE = ceramide phosphoethanolamine; PE = egg yolk phosphatidylethanolamine, Sa = d18:0 sphinganine, Cer = d18:1/18:0 ceramide. From the sixth lane onward are standard Folch (non-acidic) lipid extractions from: BT iSPT 0 ng/mL aTC induction (no SL), WT BT, *Bacteroides uniformis* (DSM 6597), *Bacteroides fragilis* (DSM 2151), *Bacteroides vulgatus* H5_1 (DSM 108228), *Bacteroides vulgatus* (DSM 1447), *Prevotella veroralis* (ATCC 33779), *Prevotella copri* (DSM 18205), *Porphyromonas gingivalis* (DSM 20709), *Flectobacillus major* (DSM 103), *Acetobacter malorum* (DSM 14337), *Sphingomonas paucimobilis* (ATCC 29837), and *Novosphingobium acidiphiulum* (DSM 19966). Homology to BT protein sequences in the inositol lipid cluster using NCBI BlastP (at e-values below 0.001) are indicated below species names with a white circle. Bacteroidetes spp. are on a blue background; alpha-Proteobacteria spp. are on a green background. (C) Predicted structures and ion chromatograms of PI-DHC structures in lipids extracted from the diverse sphingolipid-producing species shown in the same order in panel B. Phosphoinositol dihydroceramide structures include C34(OH) PI-DHC, C35(OH) PI-DHC, C36(OH) PI-DHC, C35 PI-DHC, and C36 PI-DHC.

These results indicate that inositol lipid synthesis is tied in a regulatory fashion to capsule specificity in BT. The CPS loci expressed in a BT population influence both recognition by the host adaptive immune system and bacteriophage predation ^40,41^. A role for SLs in mediating the interaction of a microbe with environmental stresses and external threats (*e*.*g*. antibiotics such as polymyxin and phages, respectively) has also been shown in the freshwater bacterium *Caulobacter crescentus*, which responds to phosphate starvation by producing complex glycosphingolipids ^42^. Given the remodeling of the capsule observed here when inositol pathways are genetically manipulated, inositol synthesis in BT could influence BT capsule detection by host immunity or phage. The regulatory link between inositol lipid synthesis and capsule specificity implies an indirect role of inositol in host-BT interactions.

We assessed how widespread the capacity for inositol lipid synthesis is in the Bacteroidetes. Inositol SLs have only been described in few bacterial species to date ^14–16^. We investigated the extent of this biosynthetic capacity across ten representative members of the phylum Bacteroidetes and in other known bacterial SL-producers. Using a homology cutoff of an e-value less than 1e-8, we compared the BT amino acid sequences for MIPS, PIPS, PIPh, PI-DHC synthase, and the BT InsP6 phosphatase, MINPP ^43^ (BT_1526, BT_1523, BT_1525, BT_1522, and BT_4744, respectively) to these related genera (Fig. 5B; Table S4). TLC analysis of lipids from these species revealed that most species with homology to the BT MIPS and PI-DHC synthase had lipid bands consistent with PI and/or PI-DHC in line with their genomically-predicted capacity (the exception was *Flectobacillus major*, a Proteobacterial species with genes encoding protein homology but that did not produce inositol lipids under tested conditions). However, we were surprised to also observe lipid bands consistent with the synthesis of PI and PI-DHC in species lacking homology to BT_1522/23/25. HPLC-MS analysis of these lipids confirmed that two species genomically predicted to lack inositol lipids (*B. vulgatus* and *P. veroralis*) in fact produced the same PI-DHC species as those with homology to BT_1522/23/25 (*B. uniformis* and *P. copri*) (Fig. 5C).

To understand this unexpected result, we searched the genomes of related species containing a BT_1526 (MIPS) homolog but lacking homology to the remainder of the BT cluster. Using PHI-BLAST with the conserved catalytic residues in BT_1523 (DX_2_DGX_2_AR…GX_3_DX_3_D) ^22^, we identified a predicted CDP-alcohol phosphatidyltransferase in the vicinity of the MIPS homolog in *Bacteroides vulgatus*. Expanding the analysis to include other genomes from the Bacteroidetes, we observed that almost every *Bacteroides/Prevotella* species containing a MIPS homolog had one of two clusters directly in the vicinity of the *MIPS* gene -either the BT-like cluster (BT_1522/23/25), or an alternate cluster including an NTP transferase (nucleotidyltransferase) domain-containing protein, CDP-alcohol-phosphatidyltransferase, and haloalkanoate dehalogenase (HAD) hydrolase (Fig. 6). The NTP transferase domain family protein (NCBI Conserved Domain Family cl11394) also shares homology with a phosphocholine cytidyltransferase motif, suggesting this protein may synthesize cytidine 5’-diphosphoinositol (CDP-inositol), similar to the synthesis of CDP-inositol as a precursor to di-*myo*-inositol phosphate solutes in hyperthermophiles ^44^. The HAD hydrolase superfamily is large and diverse, with the majority of characterized members functioning as phosphotransferases ^45^. As a lipid phosphate phosphohydrolase, this HAD hydrolase may function similarity to AUR1^46^, acting as a PI-DHC synthase like BT_1522.

**Figure 6.**
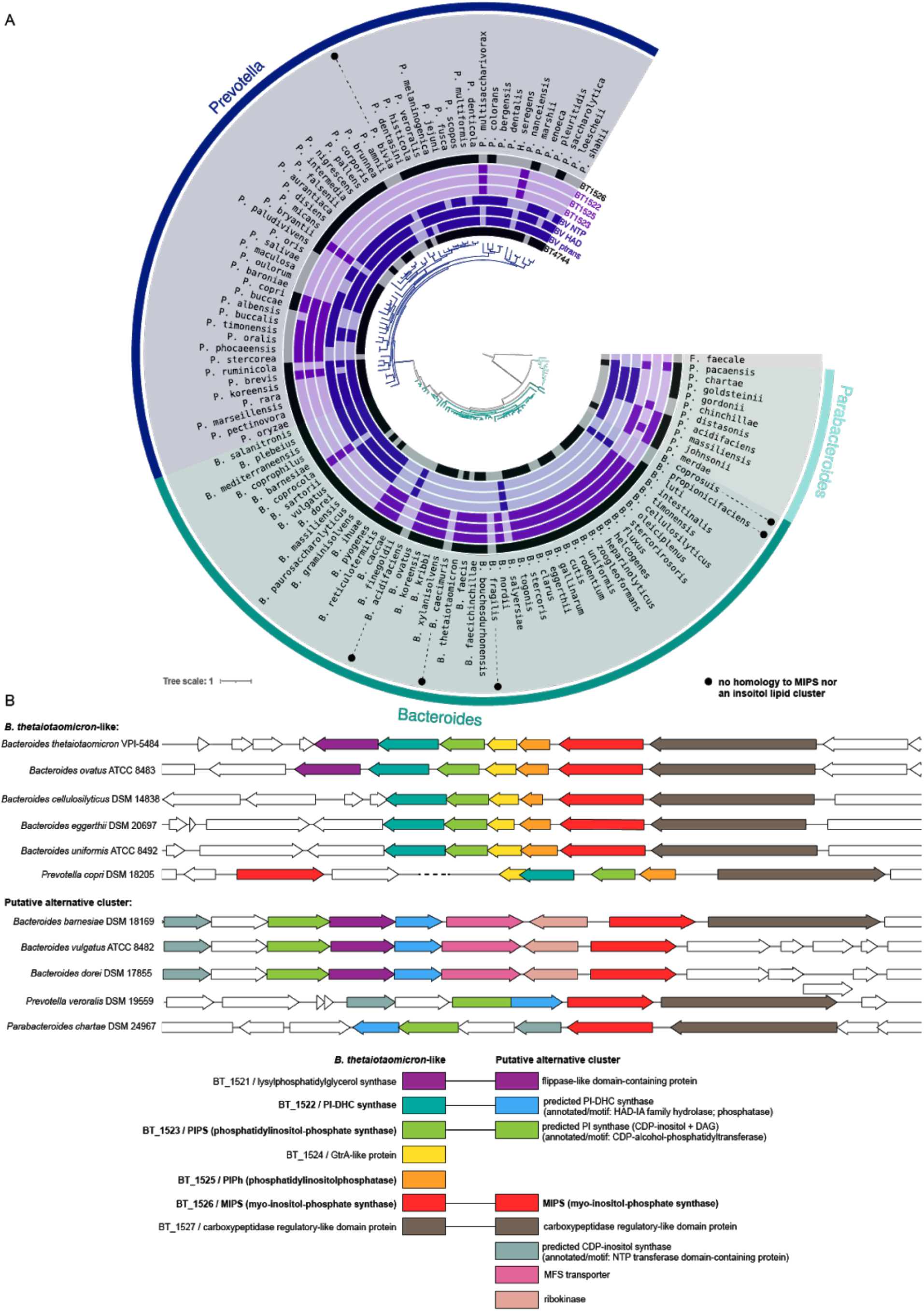
The capacity for inositol lipid synthesis is widespread within the Bacteroidetes. (A) Maximum likelihood based phylogeny of representative *Bacteroides, Prevotella*, and *Parabacteroides* species, produced from 71 conserved single copy genes present in all genomes (identified and concatenated using Anvi’o), and generated by RAxML (best tree; substitution model PROTCAT, matrix name DAYHOFF, Hill-climbing algorithm, bootstrap 50); *Flavobacterium faecale* is included as an outgroup. The rings surrounding the tree indicate species with genes that have NCBI BlastP homology to the BT inositol lipid cluster (in light purple; BT_1522, BT_1523, BT_1525, BT_1526), the BT Minpp (BT_4744), or representative proteins from the *Bacteroides vulgatus* putative alternative inositol lipid cluster (in dark purple; phosphatidyltransferase: BVU_RS13105, HAD hydrolase: BVU_RS13115, NTP transferase: BVU_RS13095). Homology at an e-value below 1e-8 is indicated by dark coloration in the inner circles. (B) Genomic regions surrounding the BT_1526/MIPS homolog in representative Bacteroidetes, compiled using the PATRIC 3.6.9 Compare Region Viewer. Protein homology (determined using NCBI-BlastP) to proteins in the BT-like inositol lipid metabolism cluster (left in key) or the *Bacteroides vulgatus*-like putative alternative inositol metabolism cluster (right in key) is indicated by color. The functions of enzymes in bold were characterized in this study; sequences with predicted redundant functions between both clusters are linked in the key.

The functions of these genes (HAD hydrolase, NTP transferase domain protein, and CDP-alcohol-phosphatidyltransferase) are not confirmed, but offer an alternative pathway enabling synthesis of PI-DHC without a PIP intermediate (similar to PI synthesis in eukaryotes ^47^), with PI synthesis resembling the synthesis of phosphatidylethanolamine or phosphatidylcholine in the Kennedy pathway ^48,49^. Following this logic, the NTP transferase protein would first synthesize CDP-inositol from *myo*-inositol phosphate and CTP. CDP-inositol and a diacylglycerol (DAG) substrate would then be converted to PI, and PI would be converted to PI-DHC by the HAD hydrolase. The MIPS homolog is most commonly clustered directly with these other genes, with some exceptions (*e*.*g*., *Prevotella copri*) (Fig. 6). Interestingly, in *P. veroralis*, the CDP-alcohol phosphatidyltransferase and HAD hydrolase proteins are fused (Supp. Fig. 5), suggesting the possibility for a cohesive single-enzyme conversion of CDP-inositol to PI-DHC through a PI intermediate. Some additional putative enzymes are shared in the vicinity of both clusters, including genes annotated as a lysylphosphatidylglycerol synthase (BT_1521 homolog) and a carboxypeptidase-regulatory-like domain protein (BT_1527 homolog) (Fig. 6). This alternative pathway could explain PI-DHC synthesis by *P. veroralis* and *B. vulgatus* despite lack of homology to the BT inositol lipid cluster (BT_1522/23/25).

Among the Proteobacteria species tested, *Sphingomonas paucimobilis* and *Novosphingobium acidiphilum* did not make hydroxylated SLs (data not shown). Despite lacking homology to either the BT-like or putative alternative inositol lipid cluster, *N. acidiphilum* produced a SL with a retention time and headgroup fragmentation consistent with *Bacteroides* PI-DHC fragmentation (Supp. Fig. 2A). *S. paucimobilis* also produced a lipid with fragmentation similar to *Bacteroides* PI-DHC, but did not produce the fragment at 241 m/z, suggesting a phosphorylated-hexose DHC unlike those produced by *Bacteroides* (Supp. Fig. 2B). In addition, the TLC analysis (Fig. 5B) shows a lipid band in the PI/PI-DHC region for mouth-associated *Porphyromonas gingivalis*, which is likely a phosphorylglycerol-DHC (Supp. Fig. 2C) ^50^.

To assess the distribution of the BT inositol lipid cluster (BT_1522/23/25) and the potential alternative pathway among the Bacteroidetes, we searched for homology in 162 representative species of these genera (*Bacteroides, Prevotella, Parabacteroides, Porphyromonas, Sphingobacterium*, and *Chlorobium* spp.) (Table S4). Most strains with a homolog to BT_1522, BT_1523, or BT_1525, in fact have homologs of all three of these enzymes, with a distribution that does not track phylogeny, supporting the lateral exchange of this full cluster among these host-associated species (Fig. 6). Roughly three-quarters of species we assessed belonging to the *Bacteroides, Prevotella*, and *Parabacteroides* genera have a MIPS/BT_1526 homolog, and most species with a MIPS homolog contain either the BT-like inositol lipid cluster or the putative alternative cluster. One notable exception is *Bacteroides fragilis*, which does not produce inositol lipids but synthesizes the bioactive glycosphingolipid α-galactosylceramide ^51,52^. The BT-like inositol lipid gene cluster is roughly two times more common than the alternative cluster among *Bacteroides* species, while the alternative cluster is about four times more common among the *Prevotella* species. Homologs to proteins in either cluster were absent or highly infrequent in the genera *Porphyromonas, Sphingobacterium*, and *Chlorobium*, with the exceptions of moderate alternative cluster homology in *Sphingobacteria*, and extensive BT_1525 homology in *Chlorobium* (Table S4), which may reflect a true phosphatidylglycerophosphatase function.

Our comparative genomic analyses revealed inositol lipid synthesis to be far more widespread in host-associated Bacteroidetes than previously thought. Although the putative alternative pathway remains to be functionally confirmed, the vast majority of species encoded one of the two pathways, with the alternative pathway more common in the *Prevotella*. The extensive prevalence of this function is in agreement with the widespread capacity in gut commensals for phytate (InsP6) degradation, which releases phosphorylated inositol derivatives. Although InsP6 phosphatase (Minpp) is rare across Bacteria (present in only 2.2% of completed genomes in EMBL-EBI in 2014), the majority of these enzymes are found in gut microbiome-affiliated species ^43^. In addition to the widespread capacity for *de novo* synthesis of inositol and its lipids reported here, these observations suggest that inositol and inositol lipid cycling in the gut are fundamental attributes of the gut microbiome.

Bacterial lipids with high structural similarity to eukaryotic bioactive lipids (*e*.*g*., SLs) have been shown to influence the metabolism and immune homeostasis of their hosts ^14,19,51,53–55^. Likewise, bacteria are already known to manipulate their host through inositol and inositol lipid metabolic pathways ^43,56^, and many bacterial and viral pathogens have also adapted to hijack the host phosphoinositide system ^57,58^. Thus, a precedent exists for trans-kingdom manipulation of inositol levels, to either a beneficial or detrimental outcome for the host. As one of the most abundant phyla within the human gut, the widespread synthesis of inositol lipids from gut-associated Bacteroidetes (*Bacteroides, Prevotella*, and *Parabacteroides* spp.) could represent a significant contribution to the lipid milieu of the gut. Of the six most prevalent and abundant *Bacteroides* species in the human gut ^59^, five have genes with homology to the BT-like inositol lipid cluster (*B. cellulosyliticus, B. eggerthii, B. ovatus*) or the potential alternative cluster (*B. dorei, B. vulgatus*), indicating potential for inositol lipid synthesis. How these lipids interact with the human host remains to be investigated.

## CONCLUSION

Inositol lipids have only recently been reported in commensal gut bacteria. In this study, we characterized the gene cluster recently hypothesized to be involved in bacterial inositol lipid synthesis in BT, in the first known study to show a functional role for these genes in the Bacteroidetes. BT synthesizes PI using a mycobacterial-like pathway with a PIP intermediate; previously, the bacterial PI synthesis pathway lacked a PIPh, which we have identified here as *BT_1525*. We also identified a putative alternative pathway for PI-DHC synthesis common among *Prevotella* species that lacks a PIP intermediate resembling the eukaryotic Kennedy pathway for phosphatidylethanolamine and phosphatidylcholine synthesis. The majority of Bacteroidetes encode one or the other of these pathways, indicating that inositol lipid production is a fundamental trait in the phylum. Together with the importance of inositol lipids in pathogen-host interactions ^60^, their high prevalence in the host-associated Bacteroidetes suggests an unexplored role in host interactions, potentially mediated both directly through provisioning and indirectly via effects on the capsule.

## Materials and Methods

### Bacterial strains and culturing conditions

Unless otherwise stated, all liquid *B. thetaiotaomicron* VPI-5482 (BT) cultures were grown anaerobically (95% N_2_ and 5% CO_2_ atmosphere) at 37°C in supplemented BHI media (BHIS; 37 g/L brain-heart infusion, 5 g/L yeast extract, 1 mg/L menadione, 1 mg/L resazurin, 10 mg/L hemin, 0.5 g/L cysteine-HCl). *E. coli* cultures were grown aerobically at 37°C in Luria broth with shaking. Final concentrations of antibiotics and selection agents were as follows: erythromycin 25 μg/mL, gentamicin 200 μg/mL, streptomycin 100 μg/mL, carbenicillin 100 μg/mL, 5-fluoro-2’-deoxyuridine 200 μg/mL. In select experiments, BT was grown in Bacteroides minimal media (BMM); per liter: 13.6 g KH_2_PO_4_, 0.875g NaCl, 1.125 g (NH_4_)_2_SO_4_, 5 g glucose, (pH to 7.2 with concentrated NaOH), 1 mL hemin (500 mg dissolved in 10 mL of 1M NaOH then diluted to final volume of 500 mL with water), 1 mL MgCl_2_ (0.1 M in water), 1 mL FeSO_4_x7H_2_O (1 mg per 10 mL of water), 1 mL vitamin K3 (1 mg/mL in absolute ethanol), 1 mL CaCl_2_ (0.8% w/v), 250 μL vitamin B12 solution (0.02 mg/mL), 0.5 g L-cysteine HCl.

For lipid analysis of non-BT strains: *Sphingomonas paucimobilis* (ATCC 29837) was grown aerobically at 30°C in nutrient broth (per L: 5.0 g peptone, 3.0 g meat extract; pH 7.0). *Bacteroides fragilis* (DSM 2151), *Porphyromonas gingivalis* (DSM 20709), *Bacteroides uniformis* (DSM 6597), *Bacteroides vulgatus* H5_1 (DSM 108228), *Bacteroides vulgatus* (DSM 1447), *Prevotella veroralis* (ATCC 33779), and *Prevotella copri* (DSM 18205) were grown anaerobically at 37°C in BHIS. *Flectobacillus major* (DSM 103) was grown at 26°C in DSM Medium 7 (per L: 1.0 g glucose, 1.0 g peptone, 1.0 g yeast extract; pH 7.0). *Acetobacter malorum* (DSM 14337) was grown at 28°C in DSM Medium 360 (per L: 5.0 g yeast extract, 3.0 g peptone, 25.0 g mannitol). *Novosphingobium acidiphilum* (DSM 19966) was grown at 28°C in DSM Medium 1199 (per L: 1.0 g glucose, 1.0 g yeast extract, 1.0 g peptone; pH 5.5).

### Generation of BT knockouts and inducible SPT strain

Genetic manipulations in the *Bacteroides thetaiotaomicron* VPI-5482 tdk (“WT”) strain were performed using double recombination from a suicide plasmid as previously described ^20^. The generation of the *BT_0870* (SPT) knockout is previously described ^54^. To create the inducible SPT (iSPT) strain, three TetR cassettes were inserted into the ΔBT_0870 genome with the constitutive PBT1311 promoter as previously described ^27^, with the native SPT (*BT_0870)* sequence reintroduced under the inducible P1TDP promoter. *BT_1522, BT_1523, BT_1525*, and *BT_1526* were knocked out using the same process in both the WT and iSPT strains. Complements for each enzyme were created in the iSPT knockout strains, likewise using the constitutive PBT1311 promoter and native BT sequences and integrated genomically, just upstream of the inositol gene cluster. Plasmids, strains, primers, and gene sequences are listed in Table S5. All constructs were verified by Sanger sequencing.

### Bacterial lipid extraction and thin layer chromatography

BT strains were grown 14-20 hours in BHIS; all other strains were grown in the media and temperatures described above to density. **“Standard” (non-acidic) lipid extraction:** Bacteria were pelleted at 3500 x g for 15 minutes, the pellet washed in PBS, and re-spun. The washed lipid pellets were lipid extracted by the Folch method ^61^, the organic fraction dried under nitrogen, the lipid film re-suspended in 2:1 (v/v) chloroform:methanol. **PIP lipid extraction:** To detect PIP, lipids were extracted according to the PI(3)P Mass ELISA Kit (Echelon Biosciences Inc.) protocol. Cells from 50-ml BHIS cultures were pelleted at 3500 x g for 15 minutes at 4°C, resuspended in 5-mL cold 0.5 M trichloroacetic acid (TCA), incubated 5 minutes on ice, and pelleted at 3500 x g for 15 minutes at 4°C. The pellets were washed twice in 3 mL 5% TCA with 1 mM EDTA, then neutral lipids were extracted twice by vortexing the pellet in 3 mL 2:1 methanol:chloroform for 10 minutes. The resulting pellets were extracted into 2.25 mL methanol:chloroform:12 N HCl (80:40:1), 0.75 mL of chloroform and 1.35 mL of 0.1 N HCl added and vortexed. The lower fraction was dried under nitrogen and resuspended in 20:9:1 chloroform:methanol:water for TLC.

### Thin layer chromatography of lipids

Lipid extracts were applied to a silica HPTLC plate with concentration zone (Supelco #60768), with loading volumes normalized to the OD_600_ of original cultures. Plates were developed in a 62:25:4 (v/v) chloroform:methanol:ammonium hydroxide system (for standard lipid extractions) or 48:40:7:5 chloroform:methanol:water:ammonium hydroxide (for PIP extractions), then sprayed with primuline (0.1 mg/mL in 4:1 v/v acetone:dH_2_O), and imaged under UV transillumination (365 nm). Lipid standards include 16:0 phosphatidylinositol (Avanti #850141), 18:1 PI(3)P (Avanti #850150), ceramide phosphorylethanolamine (Sigma-Aldrich #C4987), egg yolk phosphatidylethanolamine (Pharmacoepia), d18:1/18:0 ceramide (Cayman #19556), and d18:0 sphinganine (Avanti #860498).

### Sample Prep For HPLC-MS

Samples were frozen over liquid nitrogen and lyophilized to dryness. 1 mL of HPLC grade methanol was added to the dried material and the mixture was sonicated for 3 min (on/off pulse cycles of 2 second on, 2 seconds off, at power 100%) using a Qsonica Ultrasonic Processor (Model Q700) with a water bath cup horn adaptor (Model 431C2), with water bath flow to maintain approximately room temperature. Samples were then moved to an end-over-end rotator and extractions proceeded for 12 hours. Samples were then centrifuged at 18000 G for 30 minutes at 4 °C. The supernatant was transferred to a fresh centrifuge tube and solvent was dried with a Thermo Scientific Savant SpeedVac SPD130DLX. The dried material was resuspended in 200 μL HPLC-grade methanol, briefly sonicated, and centrifuged as before. The concentrated extract was transferred to HPLC vial with a 300 μL glass insert and stored at 4 °C until further analysis.

### HPLC-MS instrumentation

LC-MS analysis was performed on a ThermoFisher Scientific Vanquish Horizon UHPLC System coupled with a ThermoFisher Scientific TSQ Quantis Triple Quadrupole mass spectrometer equipped with a HESI ion source. All solvents and reagent for HPLC-MS were purchased as Optima LC-MS grade (Fisher Scientific).

### HPLC-MS generalized method

Mobile phase A was 94.9% water, 5% methanol, 0.1% formic acid (v/v) with 10 mM ammonium acetate. Mobile phase B was 99.9% methanol, and 0.1% formic acid (v/v). 1 µL of extract was injected and separated on a mobile phase gradient with an Agilent Technologies InfinityLab Poroshell 120 EC-C18 column (50 mm × 2.1 mm, particle size 2.7 μm, part number: 699775-902) maintained at 50 °C. A/B gradient started at 15% B for 1 min after injection and increased linearly to 100% B at 22 min and held at 100% B for 5 min, using a flow rate 0.6 mL/min. Full Scan Q1 mass spectrometer parameters: spray voltage 2.0 kV for negative mode, ion transfer tube temperature 350 °C, vaporizer temperature 350 °C; sheath, auxiliary, and spare gas 60, 15, and 2, respectively. Tandem mass spectrum analysis was carried out with Product Ion Scan mode with the following additions: collision energy: 30 V, CID gas 1.5 mTorr.

### HPLC-MS method for phosphatidylinositol phosphates

The method was slightly modified from Bui et al. 2018 ^62^. Mobile phase A was 99.9% water, 0.1% N,N-Diisopropylethylamine (v/v) with 10 uM Disodium EDTA. Mobile phase B was 99.9% acetonitrile, 0.1% N,N-Diisopropylethylamine (v/v). 3 µL of extract was injected and separated on a mobile phase gradient with an Kinetex EVO C18 UHPLC column, 2.1 × 150 mm, 1.7μm (Phenomenex, CA, PN:00F-4726-AN) maintained at 60 °C. A/B gradient started at 38% B for 6 min after injection and increased linearly to 100% B at 12 min and held at 100% B for 3 min, using a flow rate 0.35 mL/min. Full Scan Q1 mass spectrometer parameters: spray voltage 4.5 kV for negative mode, ion transfer tube temperature 325 °C, vaporizer temperature 350 °C; sheath, auxiliary, and spare gas 50, 15, and 1, respectively. Tandem mass spectrum analysis was carried out with Product Ion Scan mode with the following additions: collision energy: 30 V, CID gas 1.5 mTorr.

### Capsule monosaccharide analysis

For capsule extraction, 16-hour 20 mL BHIS cultures were normalized to OD_600_, centrifuged 3500 x g for 20 min, and gently washed two times in 50 mL of PBS (16000 x g, 4 min). The pellets were shaken (900 RPM) in 500 uL of aqueous phenol for three hours at room temperature, centrifuged at 5,000 x g for 20 min at 4°C, and the aqueous phase ethanol precipitated (cold absolute EtOH added to final concentration 80% v/v for 2 hr at -20°C, centrifuged at 18000 x g for 20 min at 4°C, washed with cold 80% EtOH and centrifugation repeated). The resulting pellet was dissolved in PBS and treated with Roth Proteinase K (1 hr at 60°C) and Merck Benzonase nuclease (20 min at 37°C). Samples were dialyzed against water (1 kDa MWCO, G-Biosciences Tube-O-DIALYZER) and stored at -80 deg C prior to inositol quantification by the UCSD GlycoAnalytics Core.

### Purification and enzymatic characterization of BT_1526

The synthetic gene encoding the BT_1526 ORF (wild type) was ordered from Genscript cloned into a pET-28a expression plasmid with a six-histidine tag at the N-terminus. The pET-BT_1526 plasmid was used to transform *E. coli* BL21 (DE3) cells for overexpression. The BT_1526 MIPS protein was expressed by culturing the transformed cells in LB medium supplemented with 35 ug/ml kanamycin at 37 °C 200 rpm shaking, until the cells reached the mid-exponential growth stage (OD_600_ = 0.5). Protein expression was then induced by the addition of 0.1 mM IPTG for five hours with a reduced temperature of 30 °C with shaking 200 rpm. Cells were harvested by centrifugation 4000 x g, 4 °C and sonicated in 10 x v/w HisA buffer (50 mM Tris-HCl, 20 mM NH4Cl, 0.2 mM DTT and 30 mM imidazole, pH 7.5) to lyse the cells. The lysate was clarified by centrifugation at 35,000 x g, 4°C and the supernatant was loaded onto a 5 ml HisTrap column (Cytiva) equilibrated with HisA buffer. Unbound proteins were washed off the column with 20 column volumes of HisA prior to elution of the tagged BT_1526 with HisB buffer (20 mM NH_4_Cl, 0.2 mM DTT and 500 mM imidazole, pH 7.5). The protein was then subjected to size-exclusion chromatography for polishing and buffer exchange. A Superdex S200 column was equilibrated with GF buffer (50 mM Tris-HCl and 150 mM NaCl) and sample added for isocratic elution over 1.2 column volumes. Fractions from size-exclusion chromatography were analyzed by 15 % SDS-PAGE and fractions containing the pure BT_1526 protein were pooled and used in subsequent experiments. The yield of recombinant BT_1526 was typically >20mg per litre of *E. coli* culture.

### Mass spectrometry analysis of purified MIPS

Samples were analysed in the positive ion mode using HPLC coupled to a Waters Synapt G2 QTOF with an electrospray ionisation source (ESI). 5-10 μL of 10 μM protein was injected onto a Phenomenex C4 3.6µ column. The conditions for the qTOF are as follows: source temperature 120 °C, back pressure 2 mbar, and sampling cone voltage 54V. The protein was eluted with a 12 minute gradient, starting at 5% acetonitrile with 0.1% formic acid to 95% acetonitrile. The resulting spectra were processed and the charge state distributions deconvoluted using MassLynx V4.1 software.

### Assay of BT_1526 for MIPS activity

The purified BT_1526 was assayed for MIPS activity was based on a method published by Barnett ^28^ (see Fig. 3A). The assay was as follows: 1 μM enzyme, 0-50 mM D-glucose-6-phosphate, 0.8 mM NAD+ for 1 hr at 25°C. The reaction was quenched with 20% TCA then 0.2M NaIO_4_ was added for an hour at room temperature. Then 1.5M Na_2_SO_3_ was added to remove excess NaIO_4_. The reagent mix was incubated for 1 hour at room temperature, then the absorbance was measured on BioTek Synergy HT plate reader at 820 nm. Values were determined with reference to inorganic phosphate standards.

### Crystallization of BT_1526

MIPS was initially screened using commercial kits (Molecular Dimensions and Hampton Research). The protein concentration was 11.4 mg/ml. The drops, composed of 0.1 ul or 0.2 ul of protein solution plus 0.1 of reservoir solution, were set up using a Mosquito crystallization robot (SPT Labtech) using the sitting drop vapor diffusion method. The plates were incubated at 20 °C and the initial hits were suitable for diffraction experiments. The condition yielding crystals that were subjected to X-ray diffraction was PACT F6 (Molecular Dimensions, 200 mM Sodium formate, 100 mM Bis Tris Propane pH 6.5, and 20 % (w/v) PEG 3350). The sample was cryoprotected with the addition of 20% PEG 400 to the reservoir solution.

### Data collection, structure solution, model building, refinement and validation of BT_1526

Diffraction data were collected at the synchrotron beamline I04 of Diamond light source (Didcot, UK) at a temperature of 100 K. The data set was integrated with XIA2 ^63^ using DIALS ^64^ and scaled with Aimless (52). The space group was confirmed with Pointless ^65^. The phase problem was solved by molecular replacement with Phaser ^66^ using PDB file 3QVT as search model. The model was refined with refmac ^67^ and manual model building with COOT ^68^. The model was validated using Coot and Molprobity ^69^. Other software used were from CCP4 cloud and the CCP4 suite ^70^. Figures were made with ChimeraX ^71^.

### Electron microscopy

Imaging was performed by the Electron Microscopy Core at the Max Planck Institute for Developmental Biology in Tübingen, Germany. For scanning electron microscopy (SEM), cells were fixed in 2.5% glutaraldehyde/4% formaldehyde in PBS for 2 hours at room temperature and mounted on poly-L-lysine-coated cover slips. Cells were post-fixed with 1% osmium tetroxide for 45 minutes on ice. Subsequently, samples were dehydrated in a graded ethanol series followed by critical point drying (Polaron) with CO_2_. Finally, the cells were sputter-coated with a 3 nm thick layer of platinum (CCU-010, Safematic) and examined with a field emission scanning electron microscope (Regulus 8230, Hitachi High Technologies) at an accelerating voltage of 3 kV.

### RNA-seq of BT at varied levels of SPT induction

Overnight cultures were used to inoculate (in duplicate) BMM media 1:2500 uninduced, or at one of five varied anhydrotetracycline (aTC) concentrations (0, 0.2, 1.0, 5.0, 100 ng/mL), and incubated at 37°C for 14 hours to an OD600 of 0.10-0.17. Cultures were spun at 3500 x g for 15 min and RNA was extracted from the bacterial pellet with QIAzol lysis reagent and the miRNeasy Mini Kit (Qiagen). rRNA was removed with the Bacterial RiboMinus Transcriptome Isolation Kit (Invitrogen) and the library prepared with the TruSeq Stranded Total RNA Library Kit (Illumina); libraries were pooled nine per lane and sequenced by HiSeq3000 (Illumina).

Quality assessment of reads was performed using FastQC pre- and post-quality filtering with bbduk (quality cutoff = 20) ^72^. Reads were aligned to the Ensembl *B. thetaiotaomicron* VPI-5482 genome with bowtie2 and assigned using htseq-count (alignment quality cutoff = 10) ^73–75^. Differential expression analysis was performed with EdgeR and limma ^76,77^: reads assigned to rRNA genes, “ambiguous,” or “no feature” were removed, lowly expressed genes were filtered, and gene expression distributions were normalized (method Trimmed Means of M values, “TMM”). Count data from BMM samples, which were in duplicate, were further normalized by Bayes moderated variance before calculation of differential expression (adjusted p-value via Benjamini-Hochberg method). Annotations were assigned from the JGI IMG database. Heatmaps were generated with pheatmap using normalized log_2_ expression values, scaled by row with Euclidean clustering.

### Phylogenies of homology to BT inositol lipid metabolic enzymes in diverse bacteria

For the smaller phylogeny of diverse sphingolipid-producers (Fig. 5D), homology to BT inositol and inositol lipid metabolism enzymes BT_1522, BT_1523, BT_1525, and BT_1526 was identified using NCBI Blast-P ^78^ to the indicated species. For the larger phylogeny of Bacteroidetes and related genera (Fig. 5), all representative species for *Bacteroides, Prevotella, Parabacteroides, Porphyromonas, Flavobacterium, Sphingobacterium*, and *Chlorobium* genera with nomenclature recognized in the LPSN ^79^ were tested for homology to BT inositol metabolism enzymes BT_1522, BT_1523, BT_1525, BT_1526, and BT_4744. For phylogenetic comparison in both trees, 71 single copy genes present in all genomes (HMM profile Bacteria_71) were identified and concatened using Anvi’o ^80^, with alignment using MUSCLE ^81^. RAxML ^82^ was used to generate a maximum likelihood tree (Protcat substitution model, Dayhoff matrix, Hill-climbing algorithm, 50 bootstrap iterations). Strain accession numbers and Blast-P results are in Table S4.

## Acknowledgements

We are grateful to Katharina Hipp and Jürgen Berger of the Electron Microscopy Core Facility at the Max Planck Institute for Developmental Biology for their expert imaging of the bacterial capsules. We would also like to thank Andrew Goodman for providing relevant strains of *B. thetaiotaomicron*. We would like to thank Diamond Light Source (Oxfordshire, UK) for beamtime (proposal mx24948) and staff of beamline I04. Both DJC and JMW would like to acknowledge the funding provided by the Biotechnology and Biological Sciences Research Council (BBSRC, grants BB/V001620/1 and BB/V00168X/1).

## Data Availability

The BT_1526 MIPS structure analyzed during the current study is available in the Protein Data Bank repository, PDB 7NWR. Transcriptomic reads, mass spectrometry files, and all unique strains generated in this study are available from the corresponding author upon request. All remaining data generated during this study are included in this published article and its supplementary information files.

## Supplementary Figures

**Supplementary Figure 1.**
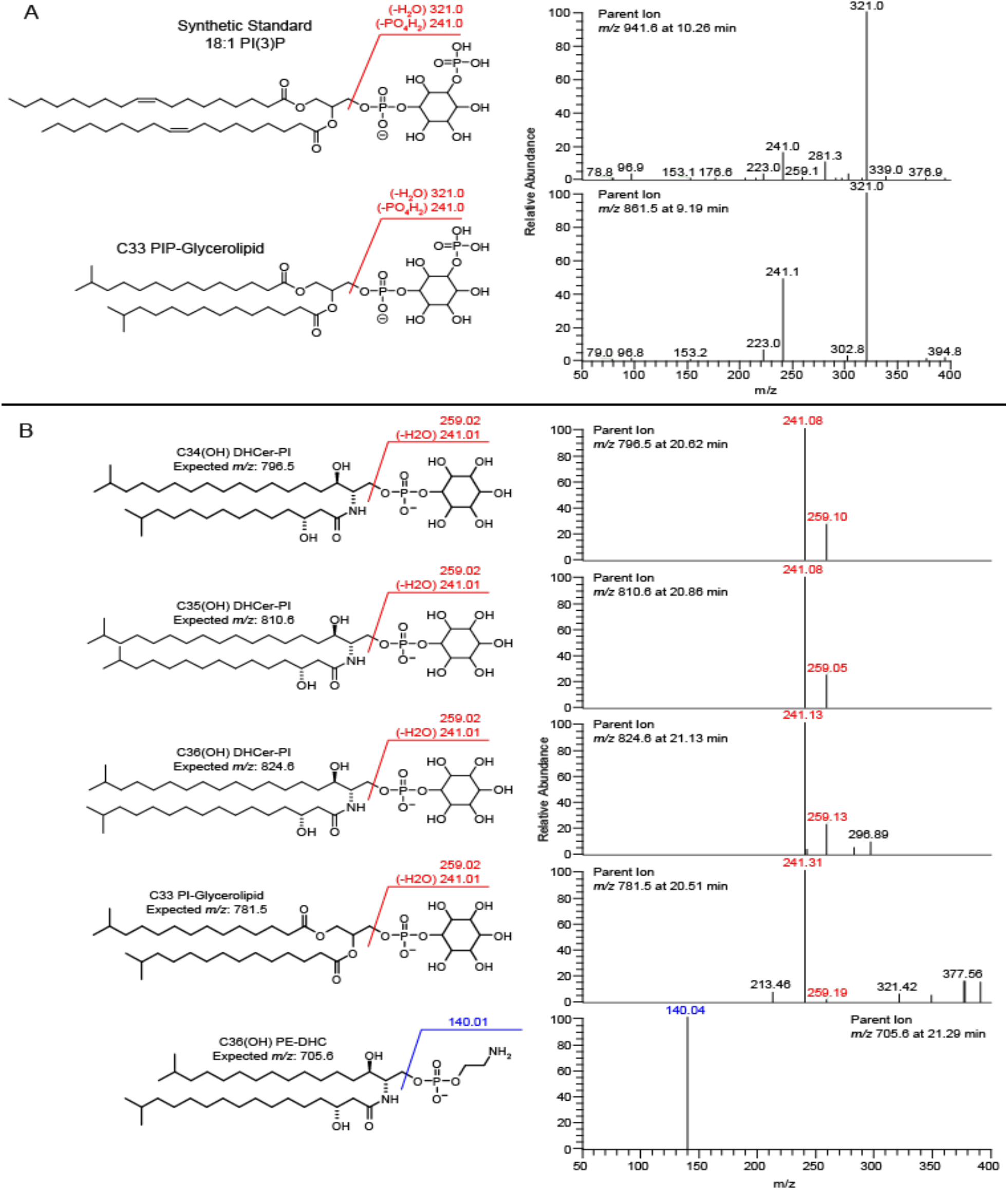
Lipid structures and fragmentation patterns of BT inositol and ethanolamine lipids. (A) Comparison of LC-MS/MS fragmentation patterns of BT-derived PIP with the synthetic standard, 18:1 PI(3)P. (B) LC-MS/MS fragmentation patterns of lipid structures present in iSPT BT at 100 ng/mL aTC induction, including PI-DHC lipids (C34(OH)DHCer-PI, C35(OH)DHCer-PI, C36(OH)DHCer-PI), C33 PI-glycerolipid, and C36(OH) PE-DHC. Loss of the phosphoinositol head group is indicated at mass 259. Fragments characteristic for lipids with phosphoinositol-based headgroups are in red; those for phosphoethanolamine-based headgroups are in blue.

**Supplementary Figure 2.**
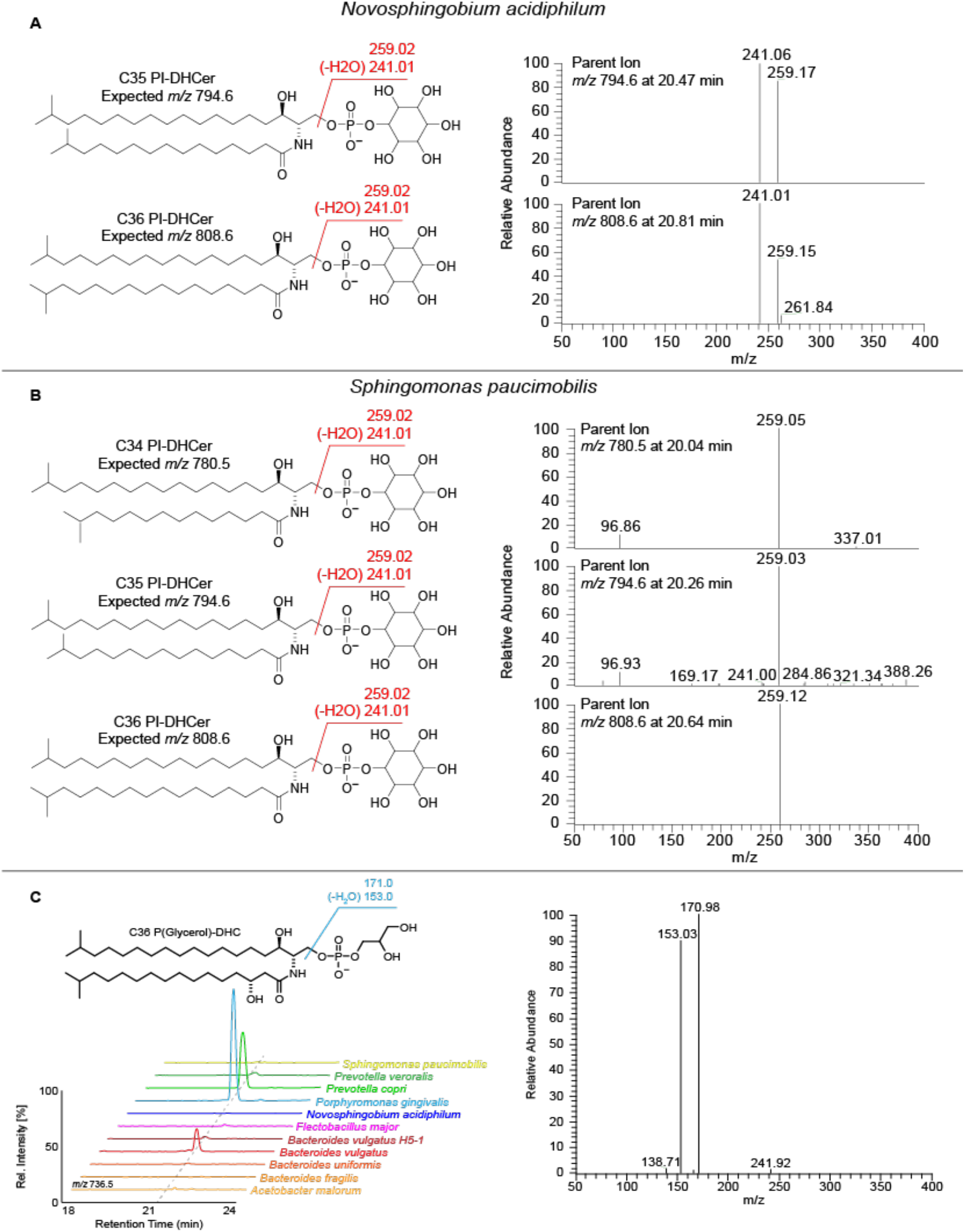
Inositol-like lipid structures in diverse sphingolipid-producing species. (A) LC-MS/MS fragmentation patterns of lipids extracted from *Novosphingobium acidiphilum* consistent with the synthesis of C35 and C36 PI-DHC. (B) LC-MS/MS fragmentation pattern of lipids extracted from *Sphingomonas paucimobilis*, demonstrating the presence of a headgroup with the same mass as inositol phosphate (259) but lacking the characteristic fragment of this group (241). (C) LC-MS/MS spectra and fragmentation pattern of a C36 P(Glycerol)-DHC structure present in *Prevotella copri, Porphyromonas gingivalis*, and *Bacteroides vulgatus*.

**Supplementary Figure 3.**
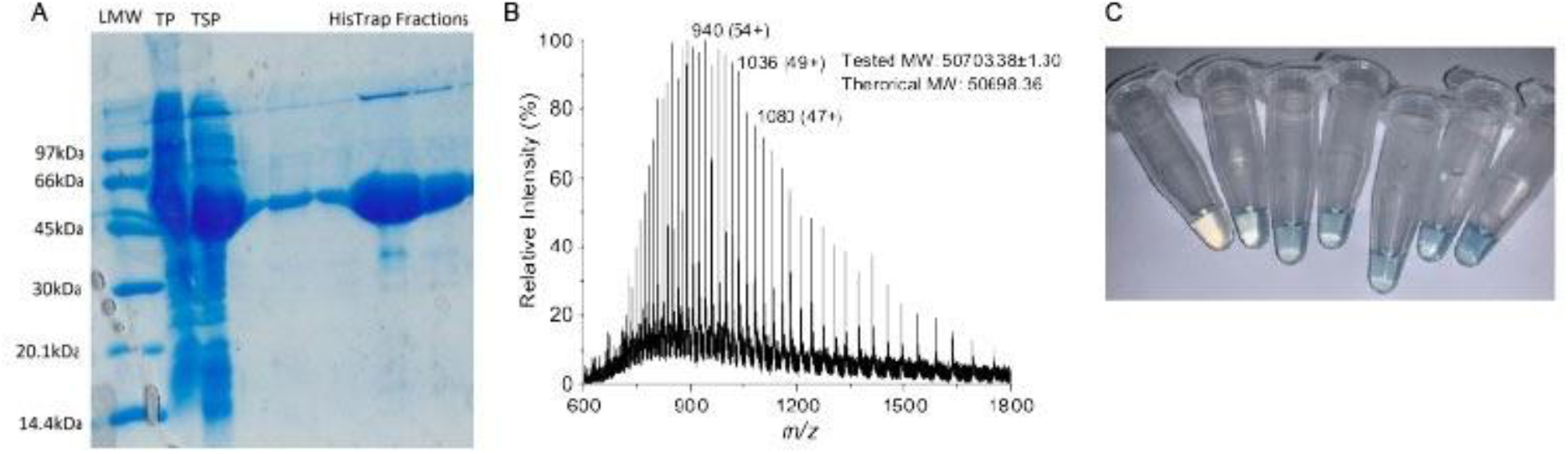
Purification of BT_1526. (A) Expression and purification of recombinant BT_1526 MIPS purified from *E. coli*. The SDS-PAGE analysis shows the purity of the samples isolated by immobilised metal affinity chromatography (IMAC). The band between the 66k and 45 kDa corresponds to BT_1526 MIPS. (B) Electrospray ionisation mass spectrometry (ESI-MS) analysis of the purified BT_1526 MIPS. Positive mode ion envelope with charge states annotated on particular masses. The predicted molecular weight matches well with the theoretical mass without the initial Met residue. (C) Typical colour observed with the molybdenum blue assay of MIPS activity at 820 nm.

**Supplementary Figure 4.**
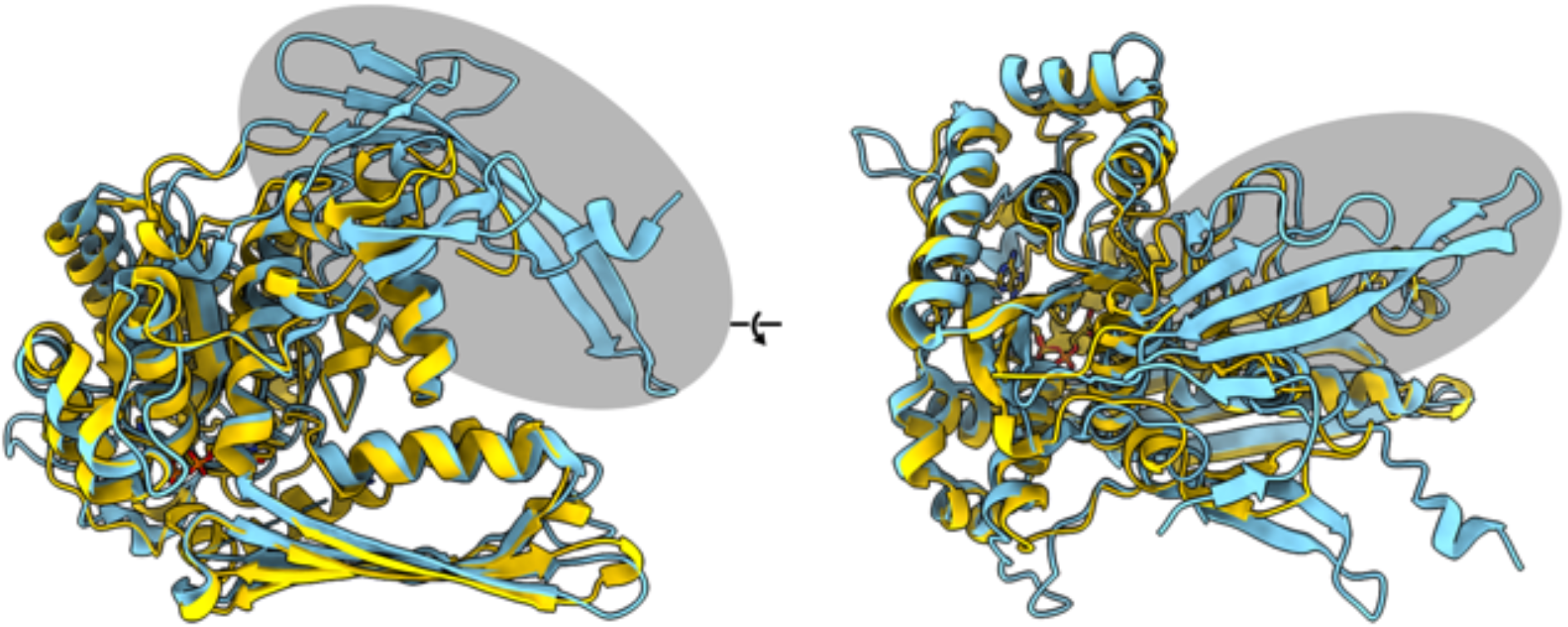
Structural comparison of prokaryotic and eukaryotic MIPS proteins. Secondary structure alignment of MIPS BT_1526 (in yellow) and *Saccharomyces cerevisiae* MIPS (PDBID: 1P1i) (in cyan). The N-terminal extension present in eukaryotic MIPS structures is highlighted with a grey oval.

**Supplementary Figure 5.**
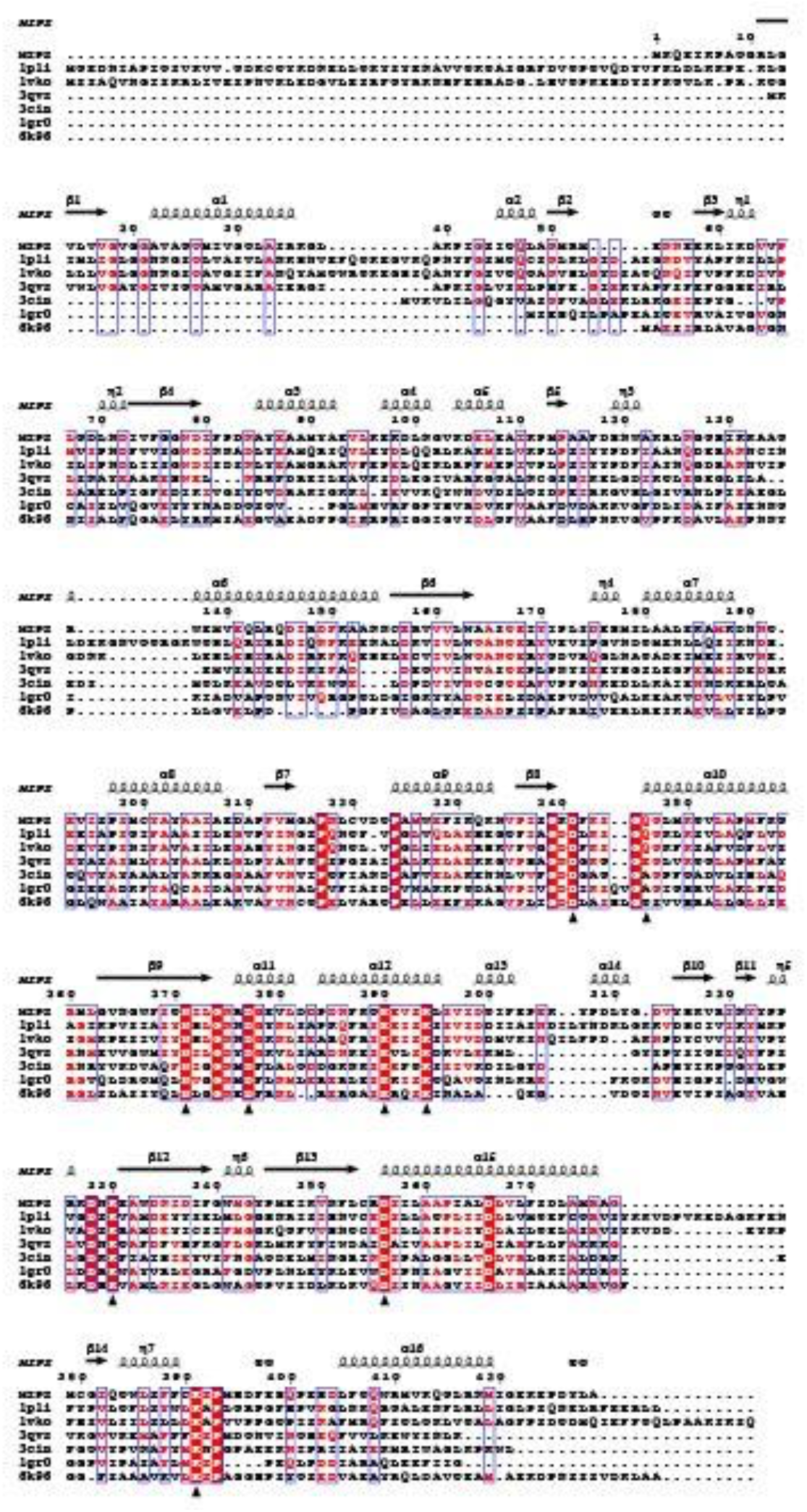
MIPS structure and sequence comparison with active site residues. The sequences are: **MIPS (BT_1526), 1p1i** (*Saccharomyces cerevisiae*), **1vko** (*Caenorhabditis elegans*), **3qvs** (*Archaeoglobus fulgidus*), **3cin** (*Thermotoga maritima* MSB8), **1gr0** (*Mycobacterium tuberculosis*) and **6k96** (*Streptomyces citricolor* Ari2). Secondary structural elements are annotated, active site residues are marked with an arrow and conserved residues marked with red shading (fully conserved) or pink shading (similar).

**Supplementary Figure 6.**
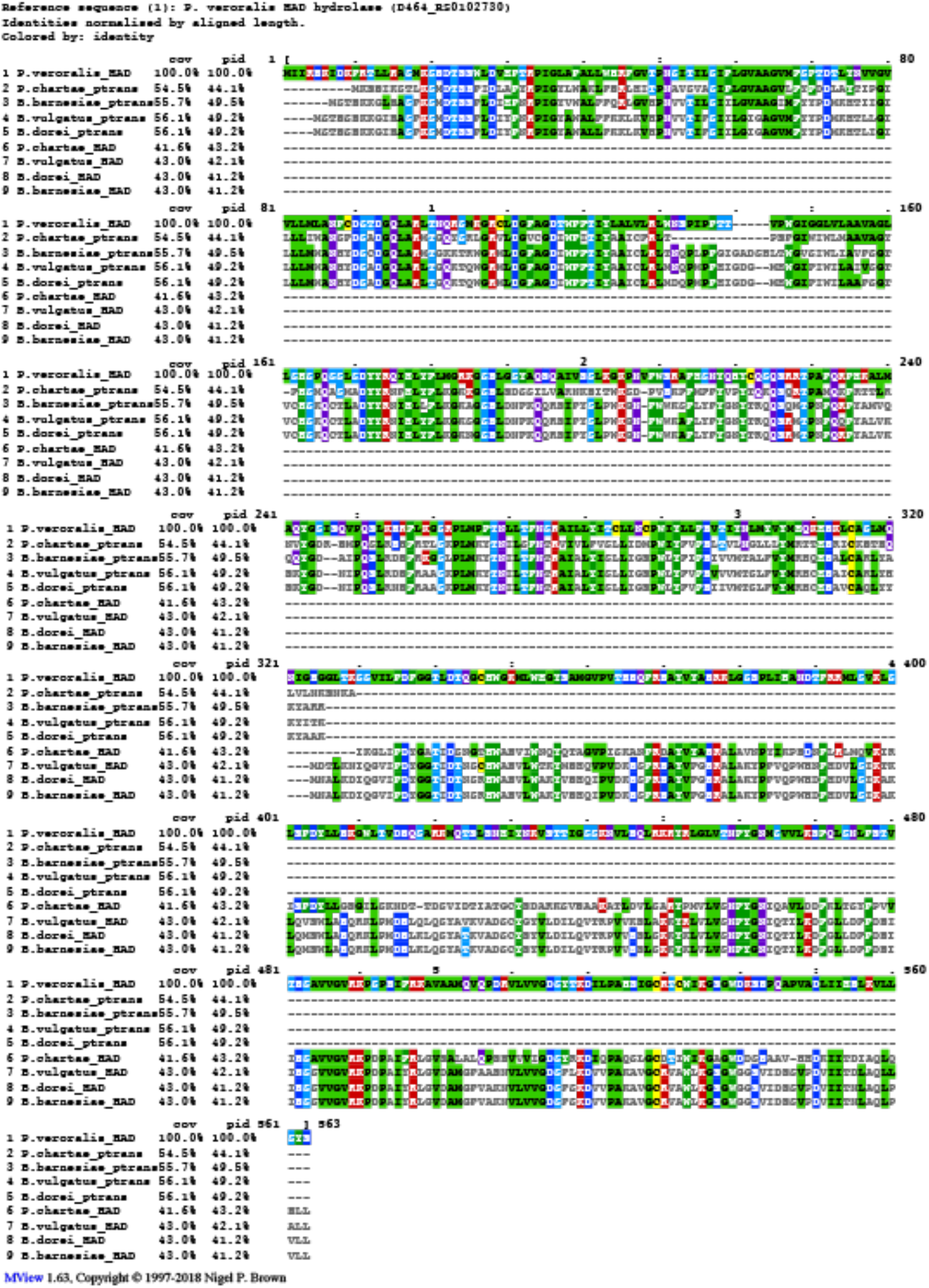
Multiple sequence alignment of predicted proteins with putative involvement in an alternative inositol lipid metabolism cluster. Amino acid sequences from representative species containing the putative alternative inositol lipid metabolism cluster (*Parabacteroides chartae, Bacteroides barnesiae, Bacteroides vulgatus*, and *Bacteroides dorei*) HAD hydrolase and CDP-alcohol phosphatidyltransferase, aligned to the *Prevotella veroralis* predicted fusion protein with homology to both of these proteins (D464_RS0102730). Alignment was performed using Clustal Omega with visualization by MView.

## Supplementary Tables

**Supplementary Table 1:** BT_1526 (MIPS) X-ray data collection and refinement statistics.

**Supplementary Table 2:** Top differentially expressed genes in wild-type *B. theta* compared to the WTΔBT_1522 strain in minimal medium with > 1.5 log_2_FC. Adjusted-P-value is Benjamini-Hochberg corrected.

**Supplementary Table 3:** Top differentially expressed genes in the iSPT strain compared to the ΔBT_1526 strain, both at 100 ng/mL aTC induction in minimal medium, with > 1.5 log_2_FC. Adjusted-P-value is Benjamini-Hochberg corrected. “CPS” column indicates the capsular polysaccharide synthesis (CPS) locus to which the gene belongs, when applicable.

**Supplementary Table 4:** E-values of BLAST-P homology to the *B. thetaiotaomicron* inositol lipid cluster, or the putative alternative pathway (using *B. vulgatus* sequences: phosphatidyltransferase BVU_RS13105, HAD hydrolase BVU_RS13115, NTP transferase BVU_RS13095).

**Supplementary Table 5:** Strains and plasmids used in this study; primers used in the amplification of genomic regions prior to plasmid assembly via restriction digest or Gibson cloning; TetR cassette components, insertion locations, primers, and gene fragment for the generation of the inducible SPT BT strain. Components and assembly were inspired by Lim et al. 2017.

**Supplementary Table 6:** Quantification of inositol in surface polysaccharides (raw values).

**Supplementary Table 7:** Kinetic data of BT_1526 MIPS (raw values).

